# The second lineage differentiation of bovine embryos fails in the absence of OCT4/POU5F1

**DOI:** 10.1101/2021.09.06.459107

**Authors:** Kilian Simmet, Mayuko Kurome, Valeri Zakhartchenko, Horst-Dieter Reichenbach, Claudia Springer, Andrea Bähr, Helmut Blum, Julia Philippou-Massier, Eckhard Wolf

**Affiliations:** Institute of Molecular Animal Breeding and Biotechnology, Gene Center and Department of Veterinary Sciences, LMU Munich, Munich; Center for Innovative Medical Models (CiMM), LMU Munich, Munich, Germany; Laboratory for Functional Genome Analysis (LAFUGA), Gene Center, LMU Munich, Munich, Germany; Bavarian State Research Center for Agriculture, Institute of Animal Breeding, Poing, Germany

## Abstract

The mammalian blastocyst undergoes two lineage segregations, i.e., formation of the trophectoderm and subsequently differentiation of the hypoblast (HB) from the inner cell mass, leaving the epiblast (EPI) the remaining pluripotent lineage. To clarify expression patterns of markers specific for these lineages in bovine embryos, we analyzed day 7, 9 and 12 blastocysts completely derived *ex vivo* by staining for OCT4, NANOG, SOX2 (EPI) and GATA6, SOX17 (HB) and identified genes specific for these developmental stages in a global transcriptomics approach. To study the role of OCT4, we generated OCT4-deficient (*OCT4* KO) embryos via somatic cell nuclear transfer or *in vitro* fertilization. *OCT4* KO embryos reached the expanded blastocyst stage by day 8 but lost of NANOG and SOX17 expression, while SOX2 and GATA6 were unaffected. Blastocysts transferred to recipient cows from day 6 to 9 expanded, but the *OCT4* KO phenotype was not rescued by the uterine environment. Exposure of *OCT4* KO embryos to exogenous FGF4 or chimeric complementation with *OCT4* intact embryos did not restore NANOG or SOX17 in OCT4-deficient cells. Our data show, that OCT4 is required cell-autonomously for the maintenance of pluripotency of the EPI and differentiation of the HB in bovine embryos.

## INTRODUCTION

During preimplantation development, the mammalian embryo undergoes two consecutive lineage differentiations resulting in a blastocyst with three distinct lineages. The trophectoderm (TE) represents the first differentiated epithelium and envelopes the inner cell mass (ICM), which retains a pluripotent state. Subsequently, within the ICM the primitive endoderm (PE), or hypoblast (HB) in human and bovine, segregates from the epiblast (EPI), which contains the last pluripotent cells and gives rise to the embryo proper. The TE will contribute the embryonic portion of the placenta and the PE/HB develops into the yolk sac [1, 2]. The fundamental mechanisms regulating these events have been studied extensively in the mouse, while advances in genome editing have enabled researchers to study the specific function of genes during preimplantation development in alternative model organisms. Given the substantial differences in regulation of lineage differentiation and maintenance of pluripotency between mouse and other mammalian species, this progress harbors the prospect of a deeper understanding of preimplantation development, also in human. Because *in vitro* embryo production techniques are highly advanced in bovine, this species offers great opportunities as a model for preimplantation development [3-5].

The second lineage differentiation, when the PE/HB and EPI segregate, is regulated by EPI precursor cells expressing FGF4, which via the MEK-pathway induces the differentiation of PE/HB precursor cells. Preimplantation embryos cultured with exogenous FGF4 develop an ICM entirely made up of PE/HB cells [4]. The transcription factor OCT4/POU5F1 plays a pivotal role in mammalian embryo development, as it regulates both the maintenance of pluripotency as well as differentiation events [6]. In mouse, loss of OCT4 prevents development of the primitive endoderm during the second lineage differentiation, while initial expression of the epiblast marker NANOG is not affected [7, 8]. On the contrary, expression of NANOG fails in OCT4-deficient bovine blastocysts, while the early presumptive hypoblast marker GATA6 is still present. Yet, it remains unclear if OCT4 has a role in the second lineage differentiation in bovine embryos, as GATA6 does not exclusively mark cells committed to the hypoblast, but also cells in the TE [9, 10].

Because data on the second lineage differentiation in bovine embryos is scarce, we first investigated expression patterns of lineage marker proteins and transcriptome dynamics of day 7, 9, and 12 embryos produced completely *in vivo* (thus representing *bona fide* samples of early bovine development). Studies of *OCT4* knockout (KO) blastocysts generated by somatic cell nuclear transfer (SCNT) and zygote injection (ZI) showed, that both EPI maintenance as well as HB differentiation is dependent on OCT4. Neither chimeric complementation with OCT4-intact blastomeres nor supplementation of exogenous FGF4 could rescue the *OCT4* KO phenotype. Therefore, we conclude that – as in mouse – OCT4 is required cell-autonomously during differentiation of the HB in bovine blastocysts.

## RESULTS

### Lineage marker and transcriptome dynamics during the second lineage differentiation in ex vivo embryos

To investigate the expression patterns of lineage markers of EPI (OCT4, NANOG, SOX2) and HB (SOX17, GATA6), we stained embryos flushed from the uterus after superstimulation as *bona fide* samples at days 7, 9 and 12 from n=7 (day 7 and 9) and n=3 (day 12) different donor cows. At day 7, OCT4 was expressed in TE and pan-ICM, and by day 9, OCT4 was restricted to EPI-cells and the percentage of OCT4 cells strongly decreased. At all examined stages, NANOG was only present in EPI cells and their precursors, i.e. co-expressed with OCT4 and SOX2 but mutually exclusive with GATA6 and SOX17. SOX2 was expressed pan-ICM at day 7 and restricted to EPI by day 9, resulting in a decreased percentage of SOX2 positive cells. Together with NANOG, SOX2 cells were still present in the embryonic disc at day 12 and it has been shown previously, that OCT4 is present in this lineage until day 17 [11, 12]. SOX17 was not present in 8/28 day 7 embryos and only faint and restricted to the ICM in the remainder. By day 9, the hypoblast began to form an inner lining of the blastocoel cavity consisting of the visceral and parietal hypoblast [13, 14], which were both marked by SOX17 until day 12. GATA6 was expressed at day 7 and 9 in the TE and ICM, but not co-expressed with NANOG. At day 12, there was no GATA6 visible in any of the lineages (Figure 1).

**Figure 1:**
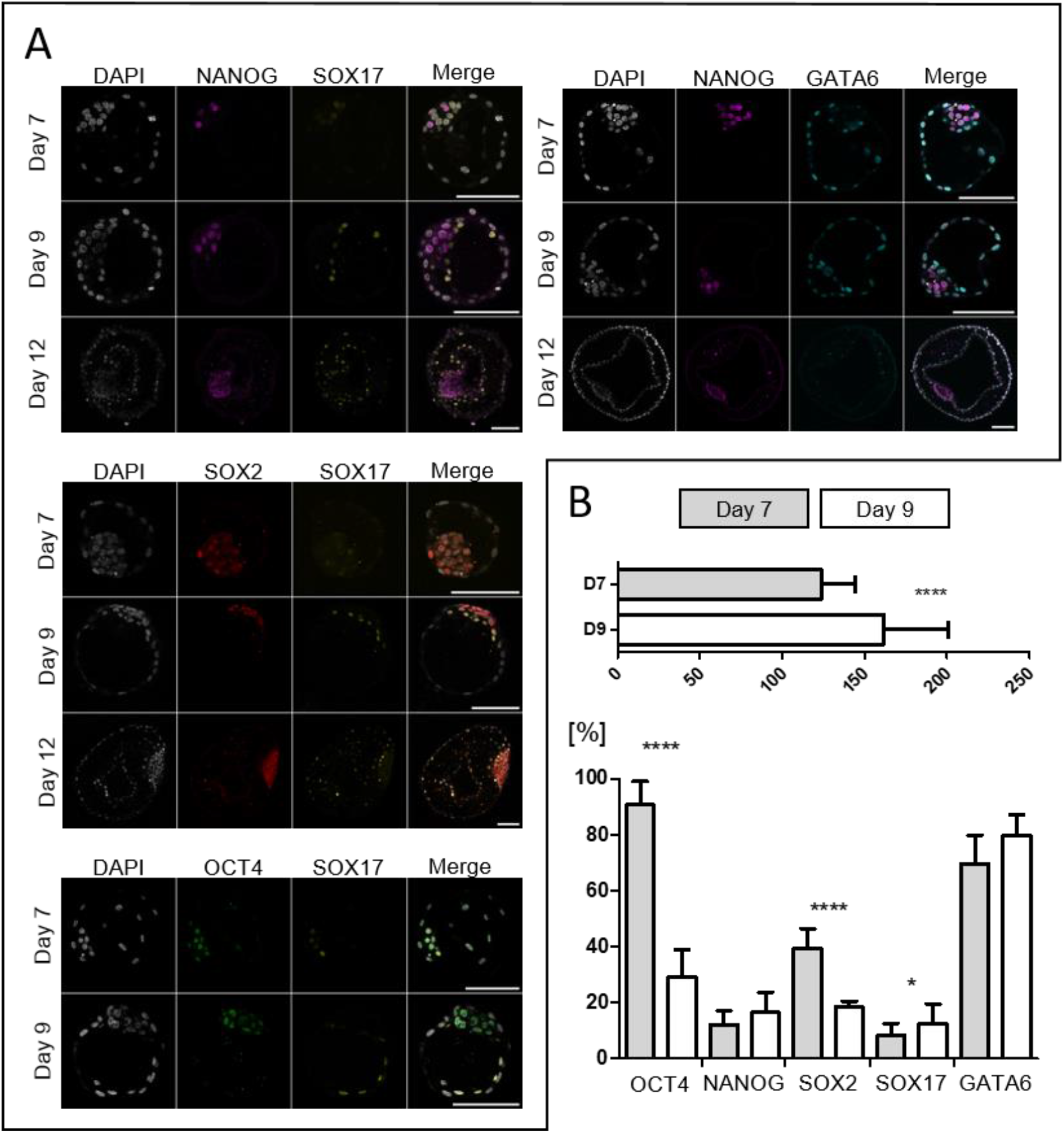
The second lineage differentiation in ex vivo embryos. A) Representative confocal planes of Day 7, 9, and 12 embryos stained for NANOG/SOX17 (n=10, 5 and 5), NANOG/GATA6 (n=10, 6 and 3), SOX2/SOX17 (n=8, 6 and 4) and OCT4/SOX17 (n=10 and 5). All scale bars represent 100 µm. B) Total cell numbers and proportion of cells stained positive for lineage specific markers at day 7 (D7) and day 9 (D9) relative to the total cell number. Data is presented as mean ± standard deviation, asterisks indicate significant differences between D7 and D9 (two tailed t-test, *P < 0.05; ****P < 0.0001). Number of examined embryos: Total cell number (D7: n=44; D9: n=21), OCT4 (D7: n=10; D9: n=5), NANOG (D7: n=17; D9: n=11), SOX2 (D7: n=15; D9: n=5), SOX17 (D7: n=20; D9: n=15), GATA6 (D7: n=10; D9: n=6). Calculation of percentage of SOX17 positive cells does not include embryos with no SOX17 expression (n=8).

In a global transcriptomics approach, we aimed to identify genes that are specific to the developmental stages at day 7, 9 and 12 and the respective embryonic cell lineages EPI, HB and TE. From three different donor cows, we analyzed three day 7 and each four day 9 and day 12 embryos. Differential gene expression analysis using DESeq2 revealed 1890 and 2716 differentially abundant transcripts (DATs, p_adj._ < 0.05) in day 9 vs. day 7 and in day 12 vs. day 9 blastocysts, respectively. DATs were categorized into eight different groups according to their gene expression pattern over the course of time, i.e., steadily increasing or decreasing, peaking at day 7, 9 or 12 and showing no difference between day 7 and 9 but increase or decrease at day 12 and vice versa. Identified DATs were compared to gene sets, which have been reported to be specific for EPI, PE/HB and TE in mouse, human and bovine embryos (Figure 2, Dataset S1). Transcripts from EPI specific genes are generally more prominent at day 7 and day 9 than at day 12; consistent with the proportion of OCT4 positive cells at day 7 and 9 (Figure 1), the abundance of *OCT4* transcripts steadily decreases until day 12. *NANOG* and *SOX2* show similar abundances at day 7 and 9 but decrease by day 12. *NODAL*, a member of the pluripotency maintaining TGFβ/ACTIVIN/NODAL signaling pathway [15], increased 80-fold from day 7 to day 9 and again decreased 2.8 fold by day 12. Interestingly, the NODAL antagonist *LEFTY2* [16] followed the same expression pattern, while transcripts of the NODAL activating convertase *FURIN* [13] steadily increased. The only EPI gene showing its highest abundance at day 12 was *FGFR1*, which in pre-gastrulation stage human embryos is reported to be enriched in hypoblast cells [17]. HB specific transcripts mostly increased until day 12, except *GATA6* and *HDAC1*. While the decreasing abundance of *GATA6* is consistent with the observed pattern in the immunofluorescence stainings, *SOX17* was not differentially abundant between day 7 and day 9 but increased later at day 12. *CDX2* is an early marker for TE, which is not differentially abundant between day 7 and day 9 but increases 1.5-fold until day 12. Except group 2 (Figure 2), TE genes are represented in every expression pattern, indicating that this lineage undergoes dynamic changes during the observed period.

**Figure 2:**
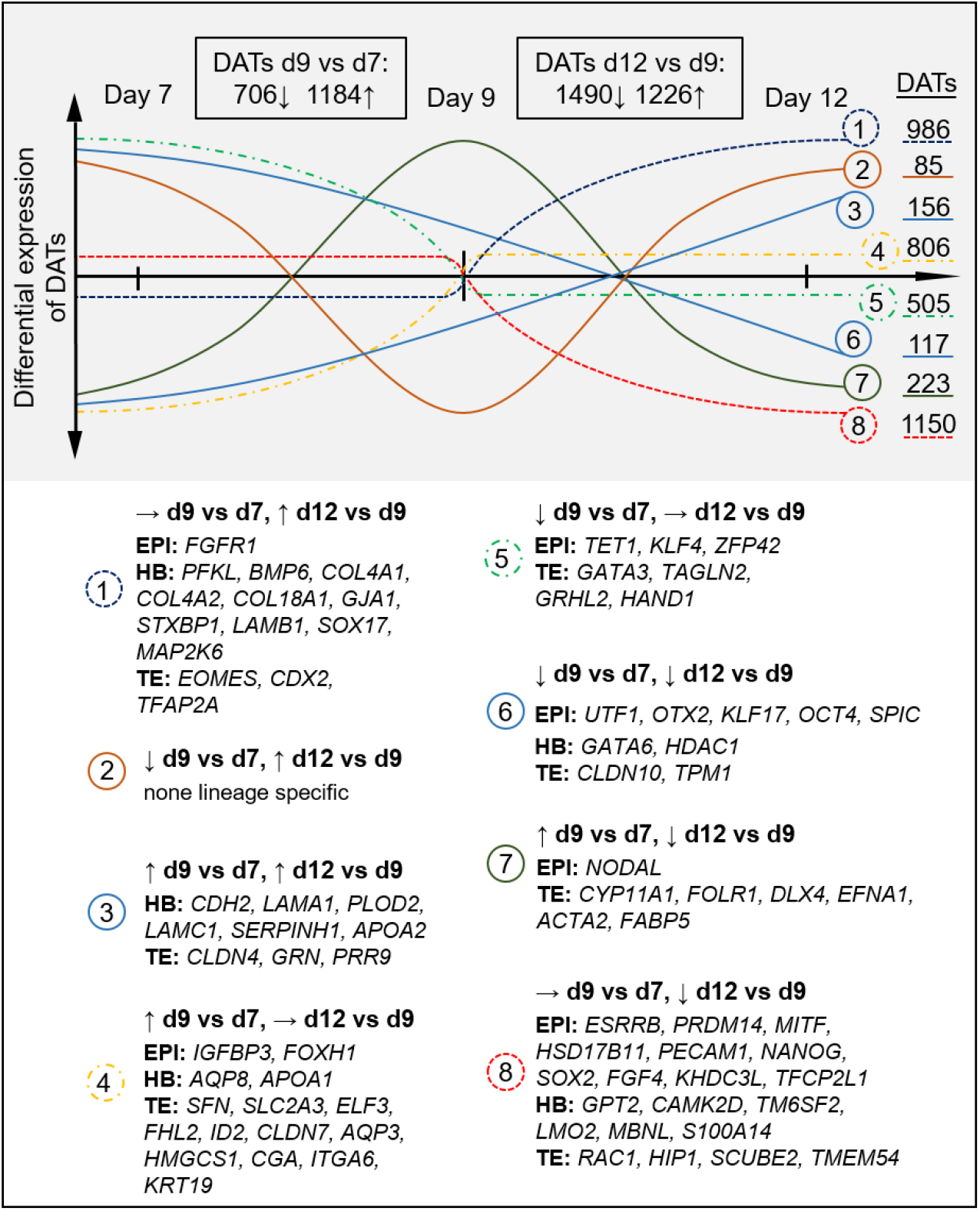
Differentially abundant transcripts (DATs) of bovine ex vivo blastocysts between day 7, 9 and 12 categorized into gene sets specific for epiblast (EPI), hypoblast (HB) and trophectoderm (TE). N=3 day 7 blastocysts and each n=4 day 9 and 12 blastocysts were analyzed using DESeq2 (padj.<0.05).

A previously published global transcriptomic dataset covering *in vitro* cultured day 7 embryos from *in vitro* fertilization (IVP Ctrl) and SCNT with wildtype cells (NT Ctrl) or cells carrying an *OCT4* KO mutation (*OCT4*KO^tm1^) [9] was reanalyzed using the current genome assembly *ARS*-UCD1.2 [18] and compared to the transcriptome profile of *ex vivo* day 7 embryos. By comparing the DATs of the three above mentioned groups against *ex vivo* day 7 embryos, we identified transcripts that were differentially abundant due to the SCNT procedure or *in vitro* culture. Five lineage specific DATs appeared in all three groups and are therefore attributable to *in vitro* culture, causing reduced abundance of *HAND1* (TE) and *HDAC1* (HB) while *HSD17B11* (EPI), *HMGCS1* and *SLC2A3* (TE) were upregulated. Two DATs were specific to the SCNT procedure with increased transcription of *CLDN7* (TE) and a lower abundance of *MAP2K6-*mRNA (HB) (Supplementary Figure S1, A). The remaining lineage specific DATs in *OCT4*KO^tm1^ against *ex vivo* day 7 contained six, two and three downregulated and one, five and eight upregulated DATs from the lineages EPI, HB and TE, respectively (Supplementary Figure S1, B), showing a shift of gene expression towards the differentiated lineages TE and HB in the absence of OCT4.

### Induction of *OCT4* knockout without targeting a known *OCT4* pseudogene

Earlier studies on the function of OCT4 in bovine embryos [9, 19] used a sgRNA-sequence, which also targets an *OCT4* pseudogene present in intron 1 of *ETF1* [20]. Therefore, we adapted a sgRNA (sgRNA2b) known to be highly efficient in human embryos [21] to the bovine orthologue sequence, where it spans an exon-intron junction at the 3’-end of exon 2 and thus does not target the pseudogene in *ETF1*, because the retrocopy does not contain intronic *OCT4* sequences [20]. The sgRNA2b sequence was cloned into PX459 V2.0 to knock out *OCT4* in somatic cells and single-cell clones were produced after selection with puromycin [22]. From n=31 single-cell clones, three retained the wildtype sequence while n=11 carried homozygous mutations, that were confirmed by a single nucleotide polymorphism (SNP) 179 bp downstream the sgRNA2b cutting site. The remaining single-cell clones had bi-allelic heterozygous (n=13) or mono-allelic (n=4) mutations. None of the single-cell clones showed any mutation at the *OCT4* pseudogene, showing that sgRNA2b specifically targets *OCT4*. From the same transfection experiment, two single-cell clones with the same homozygous deletion of two basepairs (*OCT4*^2bKOX1^ and *OCT4*^2bKOX4^) and one where no mutation had occurred (NT Ctrl^2b^) were used to reconstruct embryos via SCNT. Embryos from *OCT4*^2bKOX1^ developed to blastocysts by day 7, albeit at a much lower rate as NT Ctrl^2b^ embryos, while there was no difference between *OCT4*^2bKOX4^ and NT Ctrl^2b^ (Table 1, A). NT Ctrl^2b^ showed expression of OCT4 in both TE and ICM (n=4), while blastocysts from *OCT4*^2bKOX1^ (n=5) and *OCT4*^2bKOX4^ (n=6) stained negative. By day 8, NT Ctrl^2b^ embryos had expanded and started hatching through the incision in the zona pellucida (ZP) made during the SCNT procedure, while *OCT4*^2bKOX1^ and *OCT4*^2bKOX4^ were not able to exit the ZP and expanded to a lesser extent compared to NT Ctrl^2b^ embryos (supplementary Figure showing brightfield images will be added).

**Table 1:** Developmental rates of somatic cell nuclear transfer (SCNT) embryos

| Experimental group | *OCT4*<sup>2bKOX1</sup> | *OCT4*<sup>2bKOX4</sup> | NT Ctrl<sup>2b</sup> |
| --- | --- | --- | --- |
| No. of SCNT experiments | 7 | 4 | 9 |
| No. of fused constructs | 276 | 142 | 166 |
| Cleavage rate* (%) | 79.9 ± 11 | 83.5 ± 6.5 | 70.3 ± 15.5 |
| Morula rate* (%) | 42.7 ± 11.6 | 58.2 ± 7.7 | 47 ± 16.6 |
| Blastocyst rate* (%) | 15.1 ± 6.9<sup>a</sup> | 38.2 ± 3.7<sup>b</sup> | 40.4 ± 13.5<sup>b</sup> |

\* Data presented as mean ± standard deviation. Different superscript letters within a row indicate significant differences ( $P < 0.05$ , one-way ANOVA with Tukey multiple comparison test).

### SOX17 is lost in blastocysts lacking OCT4

To elucidate the effects of loss of OCT4 during the second lineage differentiation, we performed immunofluorescent staining of the lineage specific markers GATA6, SOX17, NANOG and SOX2 [23, 24] at day 8 blastocyst stage. In IVP Ctrl and NT Ctrl^2b^ embryos, we confirmed that at day 8 expression of SOX2 is restricted to the ICM and that NANOG and SOX17 are mutually exclusive markers of the EPI and HB, respectively. GATA6 is expressed in both ICM and TE, and GATA6 negative cells are present in the ICM. In contrast to the expression pattern in day 9 *ex vivo* embryos, SOX17 and GATA6 are always co-expressed with SOX2, indicating that SOX2 is a late EPI marker (Figure 3, Supplementary Figures S2, S3). In *OCT4*^2bKOX1^ day 8 SCNT blastocysts, there were no GATA6 negative cells and SOX2 was expressed exclusively in the ICM (Figure 3). As reported previously [9], there was no expression of NANOG and we did not detect any SOX17 positive cells (supplementary Figures S2, S3).

**Figure 3:**
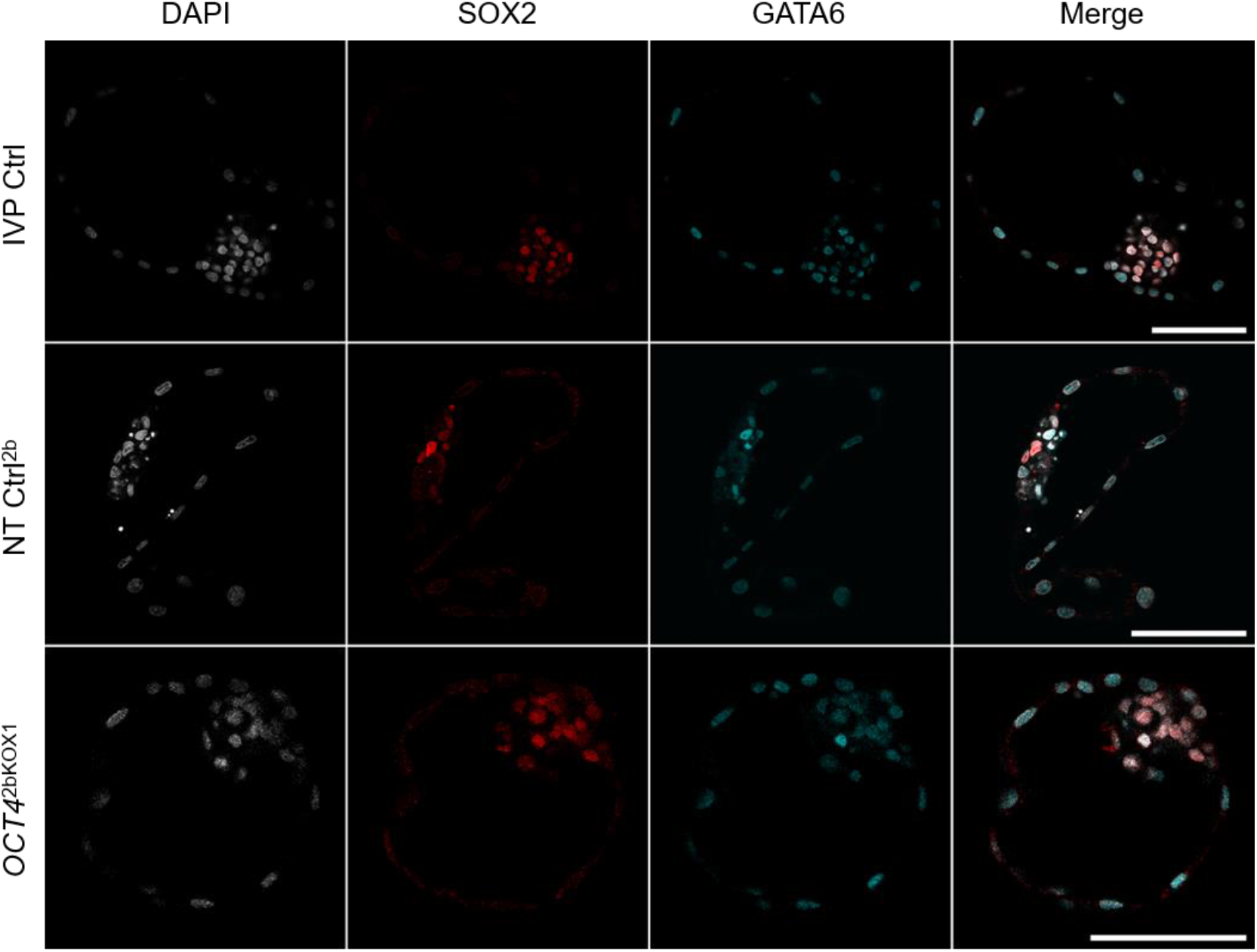
*Expression of SOX2 and GATA6 in day 8 blastocysts. Representative confocal planes of IVP Ctrl, NT Ctrl and* OCT4*^2bKOX1^ embryos stained for SOX2/GATA6 (each n=4). All scale bars represent 100 µm*.

To validate our findings from SCNT experiments, we induced KO of *OCT4* directly in zygotes from IVF by injection of a ribonucleoprotein (RNP) consisting of Cas9 protein and synthesized sgRNA2b. As control, we used an RNP with no target in the bovine genome (sgRNA Ctrl). Developmental data from 11 experiments with a total of 1224, 462 and 543 zygotes injected with *OCT4*^2bZI^, sgRNA Ctrl or non-injected, respectively, revealed that the injection procedure induced a decreased cleavage rate, but did not affect the percentage of blastocysts developed from cleaved zygotes (Table 2). To determine the mutation rate after injection, DNA was isolated from individual embryos and the targeted site was amplified for subsequent Sanger sequencing. From four experiments, we analyzed putative mutations in a total of 57 day 7 blastocysts, of which 34 had expanded. There were no significant differences in percentage of wildtype, biallelic, homozygous or monoallelic mutations between expanded and early day 7 blastocysts (Figure 4, C) and four expanded blastocysts carried homozygous mutations that induced a shift of the reading frame. Staining with antibodies against NANOG, SOX17 and OCT4 in *OCT4*^2bZI^ day 8 blastocysts in combination with genotyping after the imaging procedure enabled us to confirm absence of OCT4 on the proteome level and frame shift mutation on the genomic level in addition to the expression patterns of NANOG and SOX17. Blastocysts injected with sgRNA Ctrl showed mutually exclusive expression of NANOG and SOX17 and co-expression of both markers with OCT4 (n=8), while in *OCT4*^2bZI^ blastocysts with successful deletion of OCT4, both proteins could not be detected (n=11, Figure 4, A and B).

**Table 2:** Developmental rates of in vitro fertilized embryos after zygote injection (ZI)

| Experimental group | *OCT4*<sup>2bZI</sup> | sgRNA Ctrl | non-injected |
| --- | --- | --- | --- |
| No. of ZI experiments | 11 |  |  |
| No. of injected zygotes | 1224 | 462 | 543 |
| Cleavage rate* (%) | 66.4 ± 10.5<sup>a</sup> | 68.3 ± 12.9<sup>a</sup> | 84 ± 5.4<sup>b</sup> |
| Morula rate* (%) | 36.9 ± 10.3<sup>a</sup> | 42.9 ± 8.8<sup>a</sup> | 56.8 ± 11.1<sup>b</sup> |
| Blastocyst rate* (%) | 21.7 ± 9.9<sup>a</sup> | 27.1 ± 11.1<sup>ab</sup> | 37.2 ± 5.9<sup>b</sup> |

\* Data presented as mean ± standard deviation. Different superscript letters within a row indicate significant differences (*P* < 0.05, one-way ANOVA with Tukey multiple comparison test).

**Figure 4:**
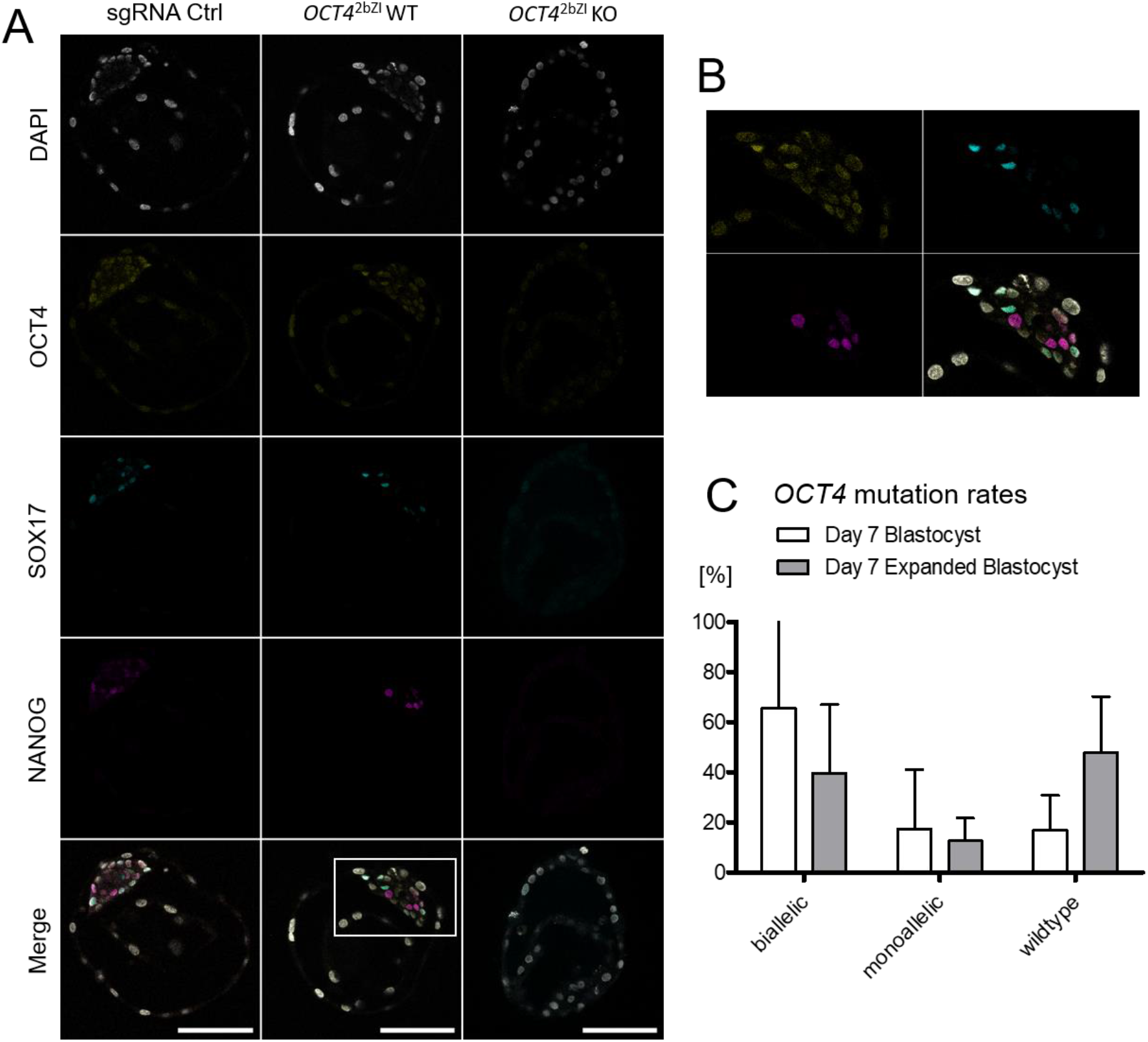
*The effect of loss of OCT4 in in vitro fertilized embryos. A) Representative confocal planes of day 8 blastocysts injected at zygote stage with a ribonucleoprotein without target (sgRNA Ctrl, n=8) or targeting* OCT4 *(sgRNA2b), where the wildtype genotype was maintained* (OCT4^*2bZI*^ *WT, n=3) or knockout induced (*OCT4^*2bZI*^ *KO, n=11). Scale bars represent 100 µm. B) Enlarged region from panel A (*OCT4^*2bZI*^ *WT merge). C)* OCT4 *mutation rates in expanded and non-expanded day 7 blastocysts (P > 0.05, two tailed t-test)*.

Developmental data from *OCT4* KO embryos produced by both SCNT and IVF show, that OCT4 is not essential for the formation of an expanded blastocyst by day 7. Expansion was present in SCNT embryos – although less pronounced – as well as in IVF embryos that carried biallelic *OCT4* frameshift mutations. Yet, a decreased blastocyst rate in *OCT4*^2bKOX1^ SCNT embryos and a higher percentage of expanded embryos where OCT4 remained intact demonstrate, that loss of OCT4 impedes the development to the expanded blastocyst stage.

### Uterine environment cannot rescue the second lineage differentiation in *OCT4* KO embryos

To evaluate if the above-mentioned phenotype of *OCT4*^2bKOX4^ embryos is alleviated or rescued when the second lineage differentiation occurs *in utero*, we transferred each four day 6 early blastocysts to five synchronized heifers and collected the embryos at day 9. As controls, we used each four IVP Ctrl blastocysts transferred to two recipients. We collected three day 9 *OCT4*^2bKOX4^ expanded blastocysts from three different recipients and five IVP Ctrl expanded blastocysts. The transferred *OCT4*^2bKOX4^ and IVP Ctrl embryos showed total cell numbers similar to *ex vivo* day 9 blastocysts (Figure 1, B) with 139.3 ± 20.5, 155.6 ± 50.28 and 161.5 ± 39.1 cells, respectively (mean ± SD, P > 0.05). Staining of NANOG and SOX17 revealed a similar expression pattern in IVF blastocysts compared to embryos completely developed *in vivo*. HB precursor cells began to form an inner lining within the blastocoel, which is confirmed by a similar proportion of SOX17 positive cells (15.1 ± 5.8 vs. 18.9 ± 4.8, mean [%] ± SD, P > 0.05), while the proportion of NANOG positive cells was markedly reduced in the IVF embryos (6.9 ± 3.8 vs. 16.5 ± 6.9, mean [%] ± SD, P < 0.05). All collected *OCT4*^2bKOX4^ blastocysts stained negative for NANOG and SOX17. Although we were not able to recover the majority of *OCT4*^2bKOX4^ blastocysts at day 9 (3/20), our data shows that in the absence of OCT4, bovine embryos survive until day 9 and expand *in utero*, but the second lineage differentiation cannot be rescued by a uterine environment (Figure 5).

**Figure 5:**
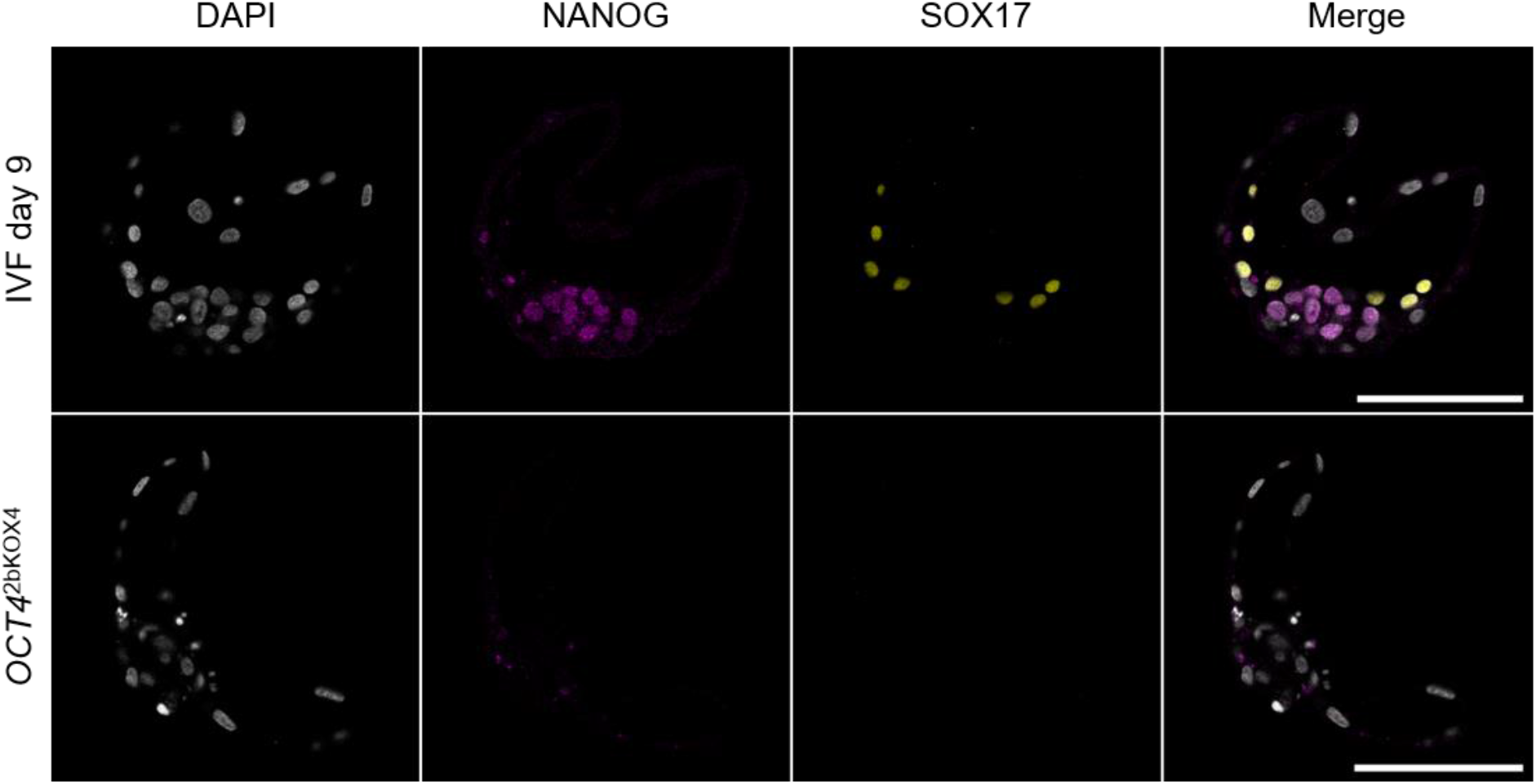
*Day 9 expanded blastocysts collected from recipient heifers stained against NANOG and SOX17. Representative confocal planes of day 9 blastocysts transferred to a recipient at day 6 and collected at day 9 from* in vitro *fertilization (IVF) or produced by SCNT using* OCT4^*2bKOX4*^ *cells. Scale bars represent 100 µm*.

### OCT4 is required cell-autonomously during the second lineage differentiation

We performed a chimera aggregation experiment in order to investigate, if OCT4 is required cell-autonomously for the expression of NANOG and SOX17. Using fetal somatic cells (FSCs), we produced a single cell clone, in which an eGFP expression vector was randomly integrated and *OCT4* was knocked out by homozygous deletion of two nucleotides in frame (*OCT4*^2bKOeGFP^). Embryos from *OCT4*^2bKOeGFP^ developed to expanded day 8 blastocysts, ubiquitously expressed eGFP and lacked expression of OCT4 (n=7), NANOG (n=8) and SOX17 (n=7). As aggregation partner, we used embryos generated from FSC wildtype cells (NT Ctrl^FSC^), which at day 8 expressed SOX17 and NANOG as expected (n=3). In three experiments, we aggregated 25 chimeras and 12 showed contribution of both *OCT4*^2bKOeGFP^ and NT Ctrl^FSC^ cells to the blastocyst. In none of these chimeras we detected co-expression of eGFP with NANOG or SOX17 (Figure 6). Therefore, we conclude that OCT4 expression in neighboring cells within the ICM cannot rescue NANOG or SOX17 expression.

**Figure 6:**
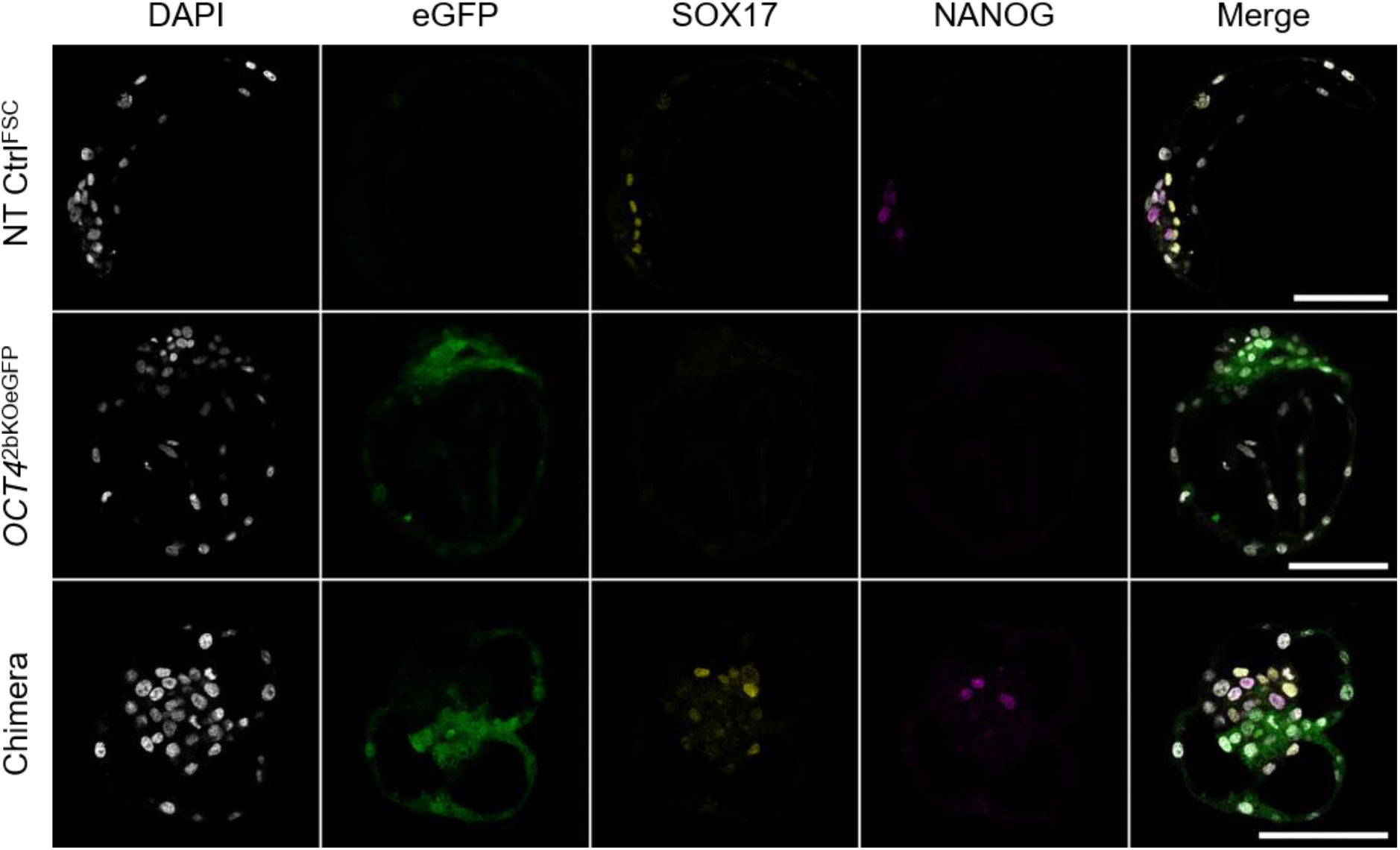
*Chimera from wildtype and* OCT4 *KO embryos. Representative confocal planes of day 8 blastocysts from SCNT using wildtype cells (NT Ctrl*^*FSC*^*) and cells tagged with eGFP carrying an* OCT4 *KO mutation (*OCT4^*2bKOeGFP*^*). Lower row in the panel shows chimera of the former embryos at day 8. Scale bars represent 100 µm*.

To further elucidate the role of OCT4 in the differentiation of the HB, we incubated *OCT4*^2bKOX4^ and NT Ctrl^2b^ with exogenous FGF4, which induces pan-ICM expression of HB-markers and ablates the expression of NANOG in wildtype embryos [25]. NT Ctrl^2b^ day 8 blastocysts showed full expression of SOX17 and no NANOG (n=10), while in *OCT4*^2bKOX4^ 10 out of 16 blastocysts showed no expression of NANOG or SOX17 (Figure 7, A). In two blastocysts, we found ectopic SOX17 expression in the TE and four blastocysts had positive cells in the ICM, albeit at a significantly lower proportion to the total cell number as FGF4 treated NT Ctrl^2b^ blastocysts (5 ± 2.4 vs. 16.3 ± 7.6, mean [%] ± SD, P < 0.05) and with a lower intensity of the fluorescent signal (supplementary Figure S4). Pairwise comparisons of the total cell number revealed a significant reduction due to loss of OCT4, which was alleviated by exogenous FGF4, while in NT Ctrl^2b^ blastocysts, FGF4 had a detrimental effect on the total cell number (Figure 7, B). Because neither chimeric complementation nor treatment with exogenous FGF4 can restore a failing differentiation of the HB in cells without functional OCT4, we conclude that OCT4 is required cell-autonomously to induce HB formation.

**Figure 7:**
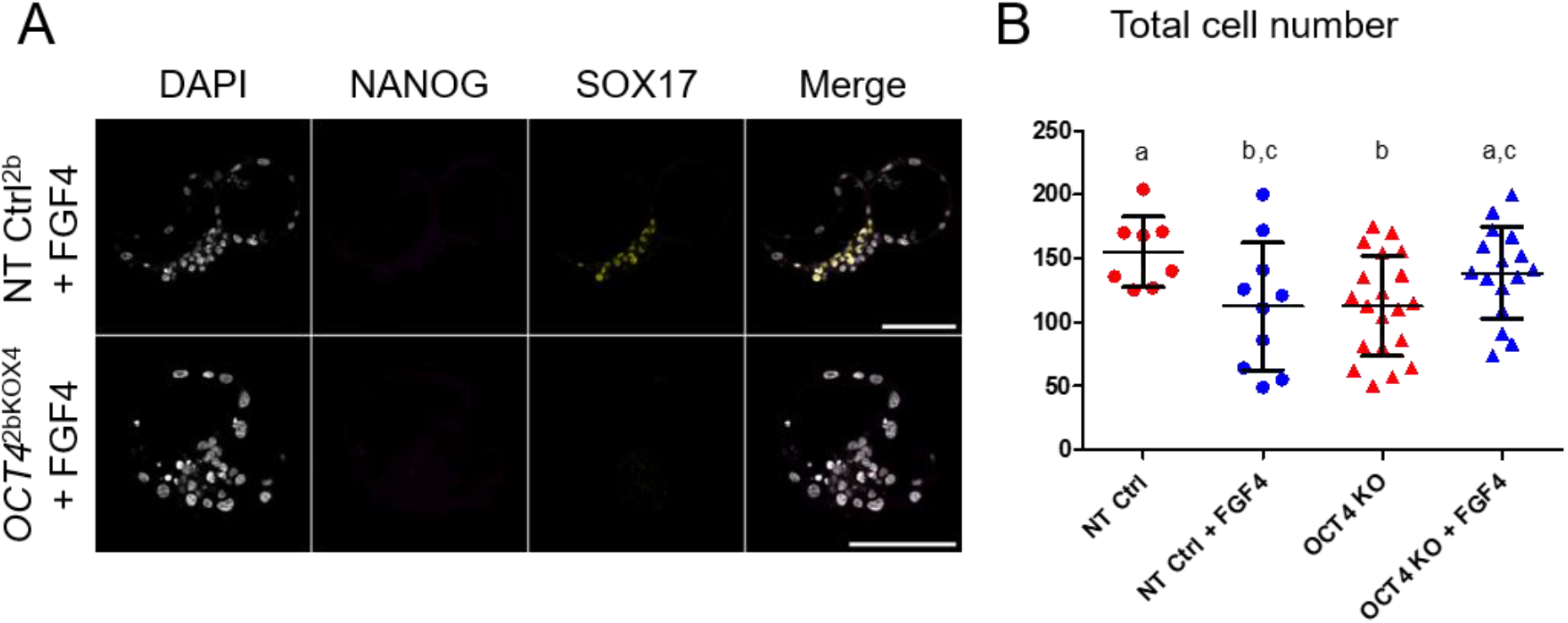
*The effect of exogenous FGF4 on* OCT4 *KO embryos. A) Representative confocal planes of NT Ctrl*^*2b*^ *and* OCT4^*2bKOX4*^ *day 8 blastocysts treated with FGF4 and heparin, stained for NANOG/SOX17 (each n = 10). All scale bars represent 100 µm. B) Total cell numbers of NT Ctrl*^*2b*^ *and* OCT4^*2bKOX4*^ *with or without FGF4 treatment. Different superscript letters indicate significant differences (P < 0.05, two tailed t-test)*.

## DISCUSSION

In this study, we set out to further elucidate the role of OCT4 in bovine preimplantation development, especially during the second lineage differentiation. Because it has not been entirely clear how the expression of known markers of the different lineages evolves during establishment of the HB-lineage, we examined the patterns in *ex vivo* derived embryos at the blastocyst stage (day 7), the expanded blastocyst stage (day 9) and the ovoid blastocyst stage (day 12). Complete HB migration by day 11 has been documented before by staining of SOX17 [26]. We were able to show, that said migration begins at day 9 with an increase of SOX17 cells and their organization into the visceral HB, ending the salt and pepper distribution of HB- and EPI-precursor cells within the ICM. Interestingly, despite an increase in SOX17 cell numbers between day 7 and day 9, we did not detect differences in *SOX17* transcript abundance. At day 7, we found a subset of embryos where SOX17 was not present yet, while in embryos that expressed the marker, intensity was low, mutually exclusive with NANOG and restricted to the ICM. Other studies using *in vitro* produced embryos report co-expression with NANOG as early as the 16-32 cell stage [10] and ectopic expression in the TE until day 6.5 [27]. In contrast to various reports using *in vitro* produced embryos [9, 10, 25], we did not find co-expression of GATA6 and NANOG in day 7 *ex vivo* embryos, indicating an earlier commitment to the HB or EPI lineage by reciprocal repression. The pluripotency factors OCT4 and SOX2 are restricted to the EPI by day 9, while at day 7 they are expressed throughout the blastocyst or the ICM, respectively, confirming SOX2 as a reliable marker for the ICM at day 7 [24]. The expression patterns described here may serve as a benchmark for assessing the quality of bovine embryos from long-term culture systems [26]. *In vitro* produced controls, that were transferred to recipients at day 6 and flushed at day 9, displayed the same total cell number and SOX17/NANOG expression pattern as completely *ex vivo* derived embryos, showing that short-term incubation *in vivo* is sufficient for stage-adequate development of the embryo, as reported previously [28].

Consistent with the staining pattern and in line with a previous report, we observed a steady reduction of *OCT4* transcripts from day 7 to day 12, while the abundance of *NANOG* and *SOX2* transcripts maintained the same levels between day 7 and day 9 and eventually decreased by day 12 [29]. Similar to human pregastrulation development, we found a decreasing abundance of transcripts associated with naïve pluripotency (*KLF4, KLF17, PRDM8, TFCP2L1, ZFP42, UTF1*) while markers for primed pluripotency, that increased in human (*FGF2, DNMT3B, SOX11, SFRP2, SALL2*), did not change from day 7 until day 12 [17]. Van Leeuwen et al. [13] detected *NODAL* transcripts in the EPI of Rauber’s layer (RL) stage (day 10-11) embryos and suggested NODAL activation through the convertase FURIN, which they detected in the RL. We found a massive increase (80-fold) in *NODAL* transcripts between day 7 and 9 together with an increase of *FURIN* and *LEFTY2*, indicating that the NODAL/BMP/WNT pathway, that later regulates patterning [30], is already active by day 9.

Studies on the effects of *in vitro* culture on the transcriptome of bovine blastocysts have mainly identified pathways related to “energetic metabolism, extracellular matrix remodelling and inflammatory signaling” [31]. While we found a total of 463 DATs between *ex vivo* and *in vitro* produced day 7 embryos, only five DATs were lineage specific, indicating that the *in vitro* culture system has no substantial effect on the mechanisms of lineage differentiation. The fact that we only found two lineage specific DATs between NT Ctrl and *ex vivo* embryos strengthens the use of embryos from SCNT to study the basic mechanisms of early lineage differentiation.

By generating *OCT4* KO embryos with a sgRNA that exclusively targets *OCT4* using both SCNT and ZI, we aimed to dissolve existing conflicts regarding the *OCT4* KO phenotype in bovine embryos. We confirmed that, regardless of the applied method, OCT4 is not essential for expansion of the blastocyst and showed that *OCT4* KO embryos survive until day 9 when transferred to a recipient cow. Daigneault et al. [19] reported, that targeting *OCT4* using ZI prevented development to the expanded blastocyst stage, while a previous report from our laboratory showed an unchanged morphology of *OCT4* KO blastocysts [9]. These two studies used the same sgRNA sequence, which also targets the pseudogene present in *ETF1*, therefore it is unlikely, that off-target effects caused the divergent phenotypes. Here we also applied ZI to delete OCT4 and observed expansion of the blastocysts, therefore we can exclude effects of the different procedures used to knock out *OCT4*. We can only speculate that the conflicting results are caused by variables in the ZI procedure or the *in vitro* culture system.

As reported previously [9], loss of NANOG was observed in all embryos without functional OCT4, but the pluripotency marker SOX2 was independent of OCT4, as reported previously [19] and similar to mouse [7]. Although there was no reduction in expression of the early HB marker GATA6, expression of SOX17 failed in the absence of OCT4. As expression of SOX17 is not dependent on NANOG [32], loss of SOX17 can be linked directly to the *OCT4* KO phenotype. In the mouse, *Oct4* KO prevents the differentiation of the primitive endoderm, not only because FGF4-MEK signaling is reduced [33] but also because OCT4 is required cell-autonomously [7, 8]. Using chimeric complementation and treatment of *OCT4* KO embryos with exogenous FGF4, we were able to show that similar to mouse, development of the HB requires OCT4 not only for the production of paracrine factors, e.g., FGF4, but also for the induction of differentiation in HB precursor cells, i.e., OCT4 is required cell-autonomously. Therefore, our data shows that, in the bovine preimplantation embryo OCT4 is required during the second lineage differentiation for maintenance of pluripotency in the EPI and differentiation of the HB.

## MATERIALS AND METHODS

### Ethics statement

All animal procedures in this study were performed according to the German Animal Welfare Act and to a protocol approved by the Regierung von Oberbayern (reference number ROB-55.2-2532.Vet_02-20-73).

### Statistics

All data were analyzed with GraphPad Prism 5.04, mean values ± standard deviation (SD) are presented. Statistical tests were two-tailed unpaired *t* test for pairwise comparisons or one-way ANOVA with Tukey multiple comparison test for analyses with more experimental groups. Level of significance was set to P < 0.05.

### Superstimulation of donors, transfer and flushing of ex vivo embryos

German Simmental heifers, 18-20 months old and 350-420 kg, served as embryo donors and recipients. Superstimulation and artificial insemination (AI) was performed as described previously [34] and the embryos were collected non-surgically by flushing at day 7, 9 or 12 (day 0 = estrous) using a flushing catheter with an enlarged tip-opening. For transfer of day 6 *in vitro* produced embryos to the uterus, the estrous cycle of recipient heifers was synchronized with a progesterone-releasing intravaginal device for eight days (PRID-alpha, Ceva, Düsseldorf, Germany) and a single dose of PGF2α analogue (500 µg cloprostenol, Estrumate, Essex, Munich, Germany) at removal of the PRID. 48-72 h later, the recipients showed signs of estrous. At day 6, embryos were transferred using a standard procedure [35] and collected at day 9 as described above.

### RNA-Sequencing and Data Analysis

Generation of RNA-sequencing libraries, sequencing, and data analysis was performed as described previously [9]. Briefly, after isolation of RNA, cDNA and RNA sequencing libraries were generated using the Ovation RNA-Seq System V2 Kit (Tecan Genomics, Redwood City, California) and tagmentation technology of the Nextera XT kit (Illumina, San Diego, California), respectively. Libraries were sequenced on a HiSeq1500 machine (Illumina) and reads were mapped to the bovine reference genome *ARS*-UCD1.2 [18] with STAR RNA sequence read mapper [36]. Differential gene expression analysis was performed with DeSeq2 [37], heat map was generated from a mean centered matrix using Heatmapper [38].

### In vitro fertilization and somatic cell nuclear transfer procedures

*In vitro* fertilization and somatic cell nuclear transfer (SCNT) were performed as described previously [39]. Presumptive zygotes and activated fused complexes from SCNT were cultured in synthetic oviductal fluid supplemented with 5% estrous cow serum, 2X of basal medium eagle’s amino acids solution 50X (Merck, Darmstadt, Germany) and 1X of minimal essential medium nonessential amino acid solution 100X (Merck). For culture of embryos with exogenous FGF4, human recombinant FGF4 (R&D Systems, Minneapolis, Minnesota) and heparin (Merck) were added at each 1 µg/ml [25].

### Immunofluorescence microscopy and image analysis

Before fixation, the *zona pellucida* was removed enzymatically using pronase (Merck) [40] or mechanically for *in vitro* produced or *ex vivo* flushed embryos, respectively. Embryos from *in vitro* culture were fixed in a solution containing 2% paraformaldehyde (PFA, Merck) for 20 min at 37° C [41] and *ex vivo* flushed embryos were fixed in 4% PFA over night at 4° C. After sequential blocking for each 1 h in 5% donkey and fetal calf serum (Jackson Immunoresearch, Ely, United Kingdom) and 0.5% Triton X-100 (Merck), embryos were transferred to the first antibody solution and incubated over night at 4° C. After washing, embryos were incubated in the second antibody solution for 1 h at 37° C and subsequently mounted in Vectashield mounting medium containing 4′,6-diamidin-2-phenylindol (DAPI, Vector Laboratories, Burlingame, California) in a manner that conserves the 3D structure of the specimen [42]. The antibodies used and their dilutions are provided in supplementary Table S1. Stacks of optical sections were recorded with a Leica SP8 confocal microscope (Leica, Wetzlar, Germany) at an interval of 1 µm using water immersion HC PL APO CS2 40X 1.1 NA or HC PL APO CS2 20X 0.75 NA objectives (Leica) and a pinhole of 0.9 airy units. DAPI, eGFP, Alexa Fluor 555, Rhodamine Red™-X and Alexa Fluor 647 were excited with laser lines of 405 nm, 488 nm, and 499 nm, 573 nm and 653 nm, respectively, and detection ranges were set to 422 to 489 nm, 493 to 616 nm, 561 to 594 nm, 578 to 648 nm and 660 to 789 nm, respectively. Cell numbers were counted manually using the manual counting plug in of Icy bio-imaging software [43], figures were produced using FigureJ software [44].

### Induction of *OCT4* knockout in fibroblast cells and zygotes

For transfection of adult ear fibroblasts, the sgRNA2b (5’ ACTCACCAAAGAGAACCCCC 3’) was cloned into pSpCas9(BB)-2A-Puro (PX459) V2.0, a gift from Feng Zhang (Addgene plasmid # 62988), using the *Bbs*I cutting site [45]. Transfection and clonal expansion after selection with 2 µg/mL puromycin for 48 h was performed as described previously [46]. Generation of *OCT4* KO cells randomly tagged by eGFP was achieved by first transfecting somatic cells derived from a fetus with a crown-rump length of 9 cm (FSC) with a linearized DNA construct and subsequently inducing *OCT4* KO via lipofection using a ribonucleotide complex (RNP) containing the sgRNA2b. The linearized construct was produced by excising the CAG-eGFP-SV40pA sequence from a plasmid, generated by introducing a de novo synthesized CAG promotor and eGFP-SV40pA [47] into the pUC57-AmpR vector backbone. The RNP was produced by mixing the synthetic and modified sgRNA2b (Synthego, Redwood City, California) and TrueCut™ Cas9 Protein v2 (Thermo Fisher Scientific, Waltham, Massachusetts) at equimolar concentrations of 8 µM in 10 mM TRIS-buffer with 1 mM EDTA. Lipofection was performed in a 6-well dish using CRISPRMAX™ Cas9 Transfection Reagent (Thermo Fisher Scientific) according to the manufacturer’s instructions. After 48 h of lipofection, eGFP positive cells were sorted individually into 96-well dishes and clonal expansion as described above was performed. Screening of single cell clones for mutations at *OCT4* and *ETF1* was achieved by Sanger sequencing as described previously [9] with primers presented in supplementary Table S2. For zygote injection, RNPs with final concentrations of 2 µM sgRNA2b or sgRNA Ctrl (5’ GGTCTTCGAGAAGACCTGCG 3’) and 1 µM Cas9 in 10 mM TRIS-buffer with 0.1 mM EDTA were produced as described above. After co-incubation of sperm and cumulus-oocyte-complexes for 14 h, cumulus cells were removed by vortexing and approximately 10 pL of the RNP were injected into presumptive zygotes using a FemtoJet4i device (Eppendorf, Hamburg, Germany). After 7 days of culture, DNA was extracted by incubating the blastocysts in a buffer containing 25 mM MgCl_2_, 1 µL/mL TritonX-100 and 150 µg/mL Proteinase K (Carl Roth, Karlsruhe, Germany) at 37° C for 1 h and subsequently at 99° C for 8 min. For Sanger sequencing, a nested PCR amplification of the *OCT4* locus was performed using 2 µL of the DNA extraction buffer directly as template. For the first PCR, we ran 25 cycles with Herculase II Fusion DNA Polymerase (Agilent, Santa Clara, California) in a 25 µL reaction volume; the second PCR used 2 µL of the first reaction as template and HotStarTaq DNA Polymerase (Qiagen, Hilden, Germany) in a 20 µL reaction volume for 15 cycles. All PCRs were performed using the buffers and instructions provided by the manufacturers, primer sequences are provided in supplementary Table S2. Extraction of DNA from fixed embryos after the imaging procedure was achieved by using the QIAamp DNA Micro Kit (Qiagen) according to the manufacturer’s instructions regarding the isolation of genomic DNA from laser-microdissected tissues followed by 35 cycles of Herculase II PCR using 4 µL of template.

### Chimera aggregation

Embryos were produced via SCNT from *OCT4* KO cells tagged with eGFP (*OCT4*^2bKOeGFP^) and FSC wildtype cells (NT Ctrl^FSC^). At day 4, the ZP was removed enzymatically and each one morula from *OCT4*^2bKOeGFP^ and NT Ctrl^FSC^ were aggregated to a chimera using phytohemagglutinin (Merck) and cultured as described previously [40]. Chimera formation was confirmed by time-lapse imaging (Primo Vision, Vitrolife, Göteborg, Sweden) and by detection of both eGFP positive and negative cells in the developed blastocysts. Chimeric blastocysts were fixed and stained as described above.

## SUPPLEMENTARY INFORMATION

Dataset S1: Lineage specific genesets and DeSEQ2 analyses

Table S1: Primary and secondary antibodies used for immunofluorescence

Table S2: Primers used for genotyping

Figure S1: The effect of in vitro culture, SCNT and OCT4 KO on the transcriptome of day 7 blastocysts

Figure S2: Expression of NANOG and SOX17 in day 8 blastocysts

Figure S3: Expression of SOX2 and SOX17 in day 8 blastocysts

Figure S4: Expression of SOX17 in FGF4 treated OCT42bKOX4 day 8 blastocysts

## ACKNOWLEDGEMENTS

This work was supported by funds from the Deutsche Forschungsgemeinschaft (DFG) under grants 405453332 and TRR127 and the Bayerische Forschungsstiftung under grant AZ-1300-17.

