## Supplementary information for "The second lineage differentiation of bovine embryos fails in the absence of OCT4/POU5F1"

*Table S1: Primary and secondary antibodies used for immunofluorescence. *SOX17 was stained with Rhodamine Red™-X in specimens with eGFP.*

| **Primary antibody** | Dilution | **Secondary Antibody** | Dilution |
|---|---|---|---|
| Rabbit anti-**OCT4** (Abcam, ab181557) | 1:250 | Donkey anti-rabbit Alexa® 555 (Abcam, ab150074) | 1:800 |
| Mouse anti-**NANOG** (ThermoFisher, 14-5768-80) | 1:250 | Donkey anti-mouse Alexa 647® (Jackson Immuno Research, 715-605-150) | 1:400 |
| Rabbit anti-**SOX2** (Millipore, AB5603) | 1:1000 | Donkey anti-rabbit Alexa® 555 (Abcam, ab150074) | 1:1000 |
| Goat anti-**GATA6** (R&D Systems, AF1700) | 1:500 | Bovine anti-goat Alexa® 488 (Jackson Immuno Research, 805-545-180) | 1:1000 |
| Goat anti-**SOX17*** (R&D Systems, AF1924) | 1:100 | Bovine anti-goat Alexa® 488 (Jackson Immuno Research, 805-545-180) | 1:200 |
| | | Bovine anti-goat Rhodamine Red™-X (Jackson Immuno Research, 805-295-180) | 1:200 |

*Table S1: Primers used for genotyping. *Primers used for Sanger sequencing, **primer used for nested PCR together with OCT4 2r.*

| | |
|---|---|
| OCT4 2f | 5‘-TTGTGGGACCTTCAAAGTAATC-3‘ |
| OCT4 2r * | 5‘-CTGCAGATTCTCGTTGTTGT-3‘ |
| OCT4 4f ** | 5‘-ATCTGGTGGATGTTGCTTTCT-3‘ |
| OCT4 12f * | 5‘-TATGTTCTTACATATCCTCTGC-3‘ |
| ETF1 2f | 5‘-TTGGGTGTGAAGTGGGTTTG-3‘ |
| ETF1 2r * | 5’-CTGGGCGATGTGGCTAATTT-3’ |
| ETF1 3f * | 5’-CTATGACTTGTGTGGAGGGATG-3’ |

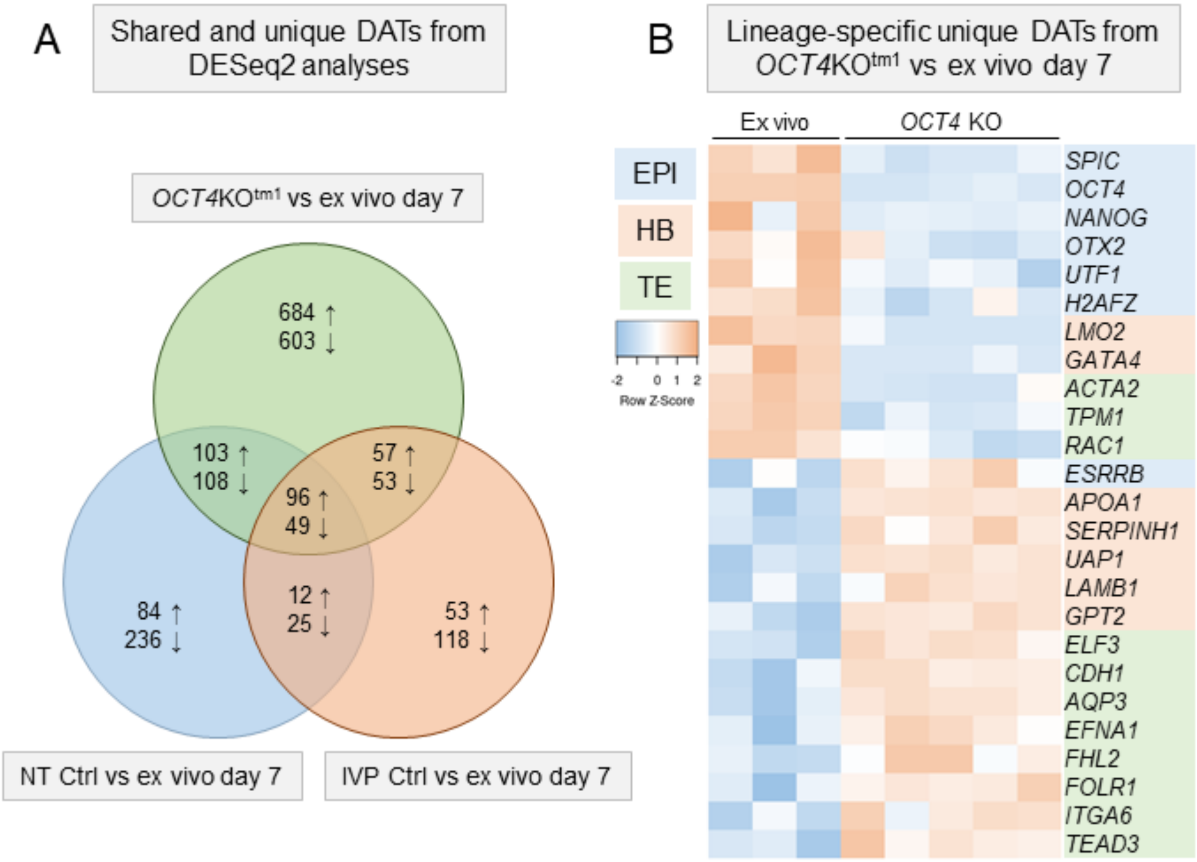


*Supplementary Figure S1: The effect of* in vitro *culture, SCNT and* OCT4 *KO on the transcriptome of day 7 blastocysts A) Venn-diagram of shared and unique differentially abundant transcripts (DATs) between day 7 blastocysts from* OCT4 *KO somatic cell nuclear transfer (SCNT,* OCT4KO*$^{tm1}$), wildtype SCNT (NT Ctrl) and* in vitro *fertilization (IVP Ctrl) against* ex vivo *embryos. B) Heat map of epiblast (EPI), hypoblast (HB) and trophectoderm (TE) specific DATs related to the loss of* OCT4 *in day 7 blastocysts. DATs were identified with DESeq2 at adjusted* P *value ($p_{adj}$) < 0.05.*

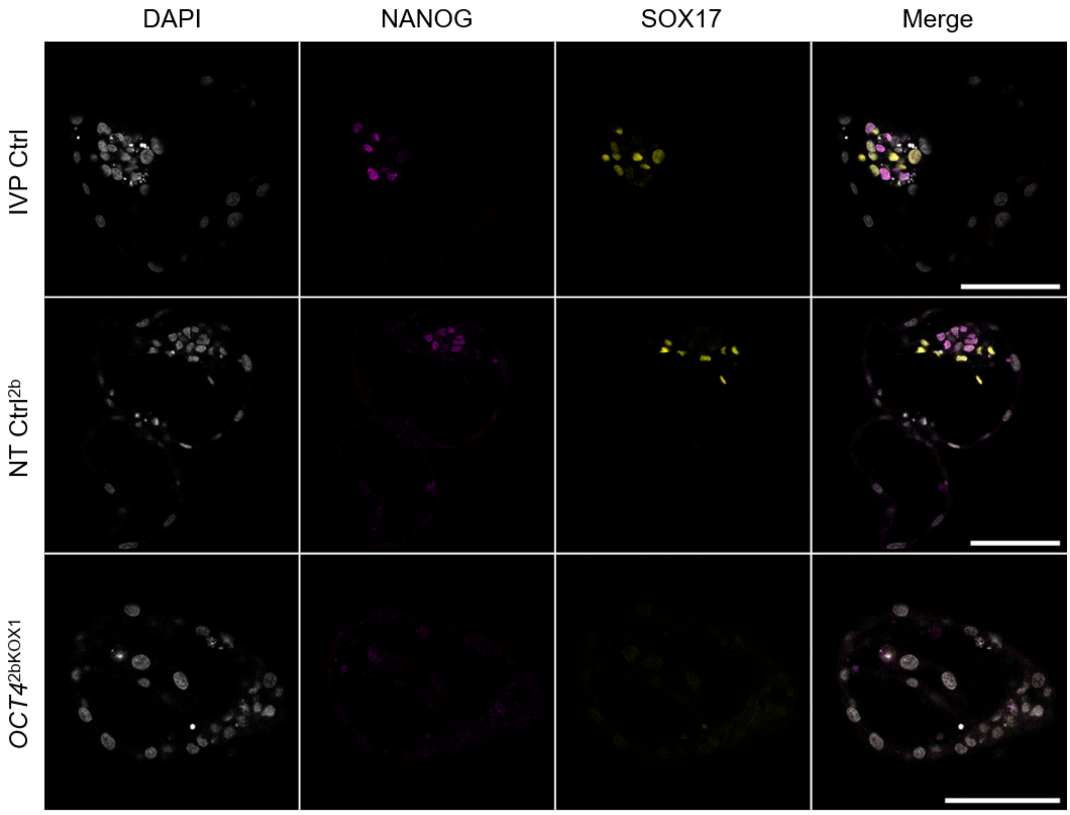


*Supplementary Figure S2: Expression of NANOG and SOX17 in day 8 blastocysts. Representative confocal planes of IVP Ctrl, NT Ctrl and* OCT4$^{2bKOX1}$ *embryos stained for NANOG/SOX17 (n = 4, 6, 14, respectively). All scale bars represent 100 µm.*

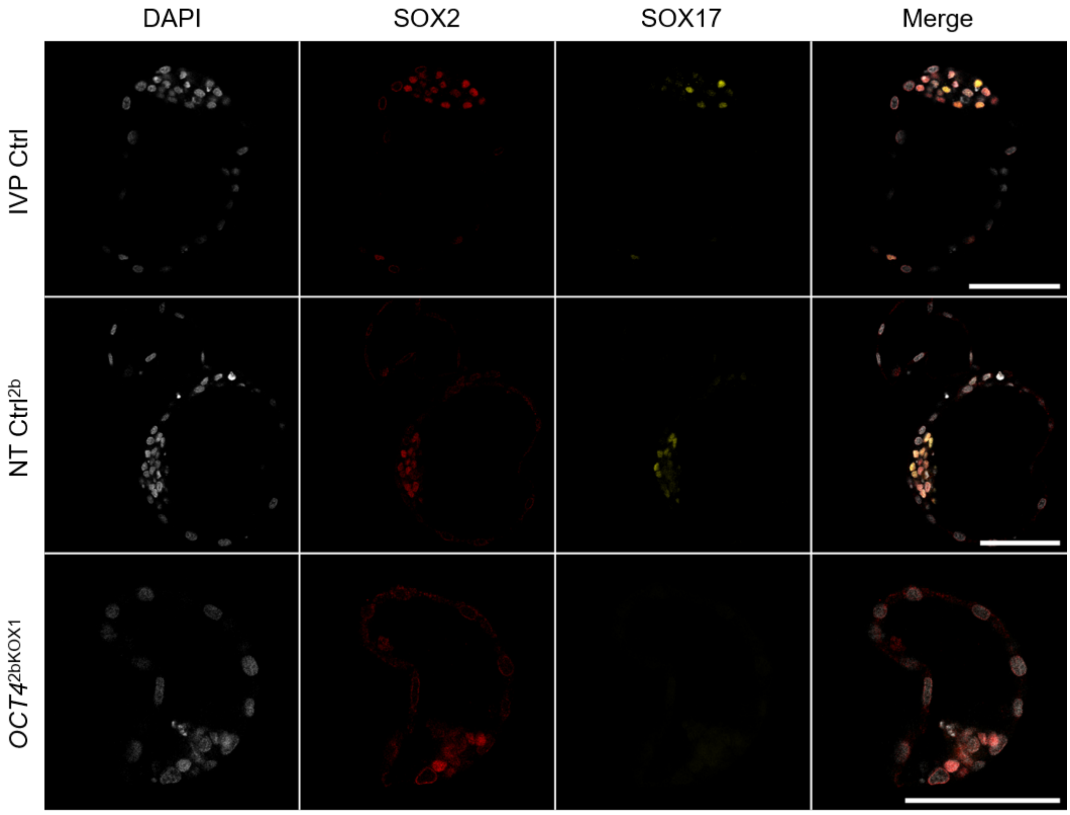


*Supplementary Figure S3: Expression of SOX2 and SOX17 in day 8 blastocysts. Representative confocal planes of IVP Ctrl, NT Ctrl and* OCT4$^{2bKOX1}$ *embryos stained for SOX2/SOX17 (n = 2, 3, 2, respectively). All scale bars represent 100 µm.*

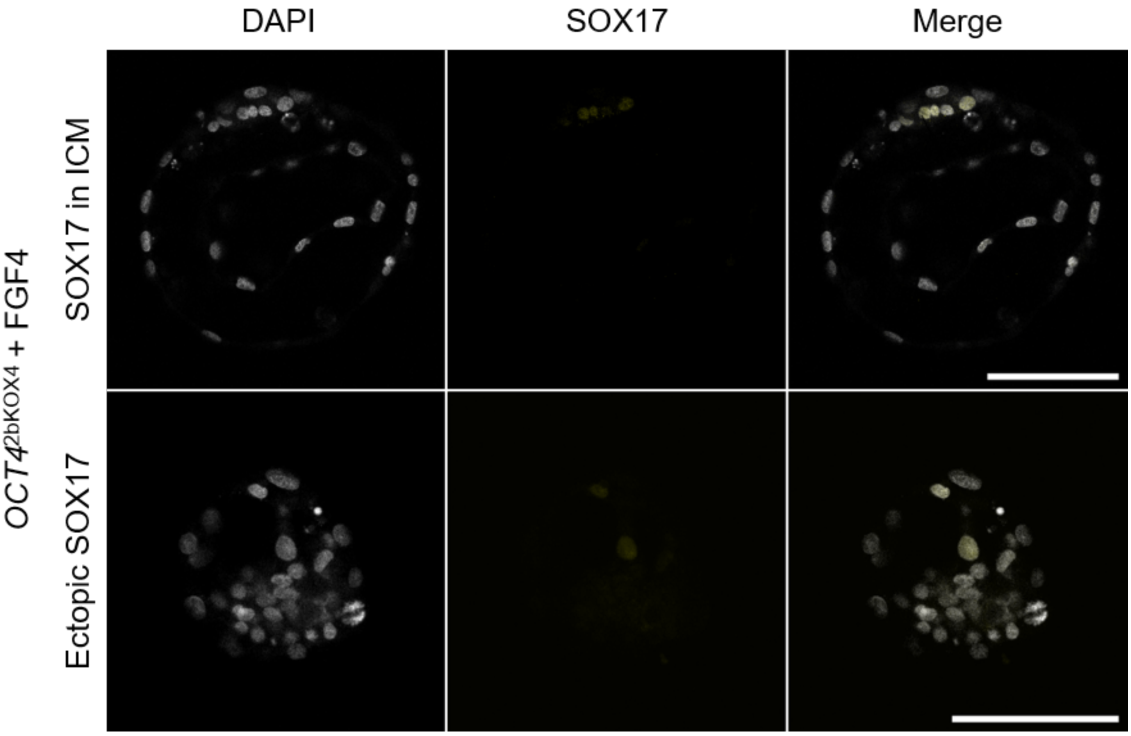


*Supplementary Figure S4: Expression of SOX17 in FGF4 treated OCT42bKOX4 day 8 blastocysts. Representative confocal planes of a blastocyst with expression in the inner cell mass (ICM, upper row, n=4) and ectopic expression in the trophectoderm (lower row, n=2). All scale bars represent 100 µm.*
